# The Cell Surface Proteome of Malignant Peripheral Nerve Sheath Tumors Reveals Therapeutic Targets

**DOI:** 10.64898/2026.03.11.711103

**Authors:** Christopher M. Stehn, Liangjun Wang, Zach Seeman, David A. Largaespada

## Abstract

Malignant peripheral nerve sheath tumors (MPNSTs) are aggressive soft tissue sarcomas and the most common cause of disease-associated death for Neurofibromatosis Type 1 (NF1) patients. In the context of NF1, MPSNTs develop from benign premalignant precursors. The transition to malignancy is usually accompanied by loss of the polycomb repressive complex 2 (PRC2), leading to aberrant upregulation of many genes. The specific mechanisms disrupted by PRC2 loss remain incompletely understood. There is a significant gap in our knowledge of which cell-surface targets become derepressed and therapeutically actionable following PRC2 loss, contributing to the current lack of effective targeted therapies for MPNSTs. This study aims to address this gap by using cell-surface capture technology with mass spectrometry to profile MPNST models. In doing so, we define PRC2-dependent effects on the cell surface proteome, including specific biological pathways that are enhanced or suppressed at the cell surface protein level. We also create an MPNST cell-surface protein compendium comprised of proteins that are highly expressed across a variety of well-defined MPNST models. We prioritized proteins that are preferentially expressed in MPNST or other cancers and for which FDA-approved therapies already exist. Specific proteins from this compendium were molecularly targeted with antibody-drug conjugates in these models to surmise their therapeutic efficacy. Results reveal PTK7 as a novel and promising target for MPNST. In total, these efforts represent a step toward addressing the knowledge gap in MPNST genesis and identifying new therapeutic targets for further testing. Additionally, this data serves as a resource for other researchers wishing to characterize specific molecular targets.

**KEY POINTS:**

PRC2 modulates key MPNST signaling pathways through the cell surface proteome
Cell surface proteomics identifies a plethora of therapeutic targets for MPNST targeted therapy
Antibody-drug conjugates targeting PTK7 show enhanced efficacy in reducing MPNST viability

**IMPORTANCE OF THE STUDY:** This study utilizes advances in biochemistry to profile the surface proteome of malignant peripheral nerve sheath tumors. In doing so, it identifies many proteins whose presence is abundant on the cell surface of MPNST cells. Pre-clinical drug testing shows that use of antibody-drug conjugates may be effective in killing MPNST cells when targeted to epitopes identified in our MPNST cell surface proteome compendium. This study is a departure from more commonly used transcriptomic methods to identify cell surface proteins by using direct surface capture and mass spectrometry, providing a more direct measurement of cell surface protein abundance. Additionally, it identifies a handful of proteins which can be directly targeted pharmaceutically and one in particular, PTK7, whose targeting is highly effective in killing MPNST cells.

## Introduction

Neurofibromatosis type 1 (NF1) syndrome is an autosomal dominant cancer predisposition syndrome caused by the inheritance of one loss-of-function copy of *NF1*, which encodes the RAS GTPase-activating protein (GAP) neurofibromin. Symptoms of this syndrome are numerous but include the development of benign and malignant tumors, pigmentary lesions, and developmental disabilities. Notable among these tumors are peripheral nerve sheath tumors (PNST)^1^. PNSTs are typically discovered as benign cutaneous or plexiform neurofibromas (cNF or pNF) and are composed of neoplastic Schwann cells (SC) that have lost the remaining *NF1* allele, along with other stromal cell types, including macrophage, mast cells, T cells, endothelial cells, and others^2,3^. The pNF lesions can become a serious concern as they may grow very large, requiring surgery or treatment with a MEK inhibitor. Importantly, they also pose a risk of ultimately progressing to a malignant peripheral nerve sheath tumor (MPNST). Patients with NF1 have a 10-13% chance of developing an MPNST within their lifetime^2,3^. Additionally, nearly half of MPNSTs present as sporadic, without an associated NF1 diagnosis, or are radiation-induced following exposure from a variety of modalities^3^. The majority of NF1-related MPNSTs develop with a well-characterized genetic etiology. Loss of heterozygosity of *NF1*, resulting in biallelic loss of *NF1* and complete loss of neurofibromin function in a Schwann lineage cell, together with local inflammatory factors, leads to the development of pNF in deep peripheral nerve sheaths^3–6^. A subset of cells in these tumors subsequently undergo loss-of-function (LOF) mutations or deletions in *CDKN2A* and often *CDKN2B.* This results in the development of an atypical neurofibromatous neoplasm of uncertain biological potential (ANNUBP), a PNST with increased nuclear atypia, hypercellularity, or other irregularities in mitotic activity^2,7,8^. The ANNUBP is thought to be the pre-malignant precursor to MPNST. Accumulation of additional mutations in ANNUBP drive formation of MPNST, including LOF mutations or deletions in *SUZ12* or *EED,* whose protein products form components of the polycomb repressor complex 2 (PRC2). Deletions in *SUZ12* or *EED* occur in 50-80% of all MPNSTs^5,9–11^. An additional feature of MPNST is an increase in genomic instability, leading to genome-wide copy number alterations and tetraploidy^12,13^.

Loss of PRC2 directly leads to loss of histone H3 lysine 27 trimethylation (H3K27me). PRC2 functions as a gene repressor during development. It controls the timing of expression for a variety of target genes through deposition of H3K27me3 at the associated gene promoter, with enrichment at CpG islands^14,15^. In the context of MPNST genesis, evidence suggests that H3K27me3 loss potentiates malignancy through increased oncogene/oncogenic pathway expression. Analysis of multiple RNAseq datasets comparing MPNST model systems with and without PRC2 activity showed that a common set of genes become upregulated following PRC2 activity loss^5,11,16^. This suggests that PRC2 typically represses a core set of genes whose increased expression contributes to malignant transformation^1^. On the other hand, some evidence indicates that PRC2 loss also results in downregulation of a small subset of genes, interferon signaling pathway. This suppression leads to an immune desert tumor microenvironment phenotype^16,17^. Finally, loss of PRC2 also increases the stemness of affected tumor cells by specific upregulation of developmental transcription programs^4^. Known direct targets of PRC2 include *HOX* loci and other transcription factors such as *FOXC1*, *HES1*, *SMO,* and *SHH*, whose dysregulation ultimately leads to a less developed phenotype^5,11^. We have identified other PRC2 derepressed drivers of MPNST, including *MET*, *PDGFRA*, and NOTCH signaling^11^. Despite these biological insights, no targeted therapies exist for MPNST, and loss of PRC2 has not been proven to be a predictive biomarker for any therapy. Furthermore, there is no consensus treatment strategy for recurrent or metastatic MPNST.

We hypothesized that cell surface proteins may represent actionable therapeutic targets for MPNST, since they could be targeted using antibody-drug conjugates (ADCs), bi-specific engagers for T-cells or natural killer (NK) cells, or chimeric antigen receptors (CARs). A 2023 study analyzed whole-cell proteomics from 23 primary MPNSTs, but it did not focus on surface proteins and could not differentiate between tumor and stromal cells^18^. Here, we have utilized a method of cell surface proteomics to identify proteins on the cell surface of models of MPNST, including cell lines and patient-derived xenografts (PDXs). Using two MPNST cell line models engineered to be isogenic except for PRC2 status, we identified surface proteins whose expression levels change in response to PRC2 activity. Additionally, we utilized a variety of tumor-derived MPNST cell lines and patient-derived xenografts (PDX) generated from MPNST tumor samples to establish a consensus MPNST surface proteome. These results provide a mechanistic understanding of the signaling changes that occur after PRC2 loss in MPNST genesis, along with a compendium of potential MPNST therapeutic targets.

## Materials and Methods

### Cell Lines and Reagents

Other MPNST cell lines and patient-derived xenografts (PDX) were kindly provided by Dr Nancy Ratner (Cincinnati Children’s Hospital) (STS-26T, S462TY, ST88-14), the Characterized Cell Line Core Facility (MD Anderson) (MPNST007), and Dr Christine Pratilas (Johns Hopkins) (JH-2-055 CL, JH-2-079c CL, JH-2-003 CL, JH-2-002 CL)^19–21^. Cell lines and media are listed in Supplementary Table S1. Antibody-drug conjugates used within the study are listed in Supplementary Table S2.

### Generation of PRC2-restored MPNST

An MPNST-derived line, S462TY, was sequentially transduced with vectors containing a reverse tetracycline-controlled transactivator (rtTA) cDNA and a tetracycline-response element (TRE) promoter-driven *SUZ12* cDNA, respectively. Following each transduction, selection was done with hygromycin at 500 ng/mL and puromycin at 5 ng/mL. Post-selection, single-cell clones were selected to obtain populations for testing restoration of H3K27me3, with optimal restoration observed under treatment of doxycycline at 500 ng/mL for 7 days. Fresh media with doxycycline was provided every two days.

### Generation of Patient-Derived Xenograft Samples

Tumors extracted from mice were minced and loaded into a Miltenyi gentleMACS^TM^ Dissociator. This process utilizes heating and cooling cycles in addition to enzymatic and mechanical separation to create a single-cell suspension. The resulting slurry is filtered and washed with PBS three times. Then, magnetic bead separation is applied to retain only the human tumor cells, creating a tumor single-cell suspension. This suspension was either frozen for future use or used directly in cell surface capture.

### Generation of Proteomic Data

Data were generated via the University of Minnesota Center for Metabolomics and Proteomics. In brief, following surface protein capture, samples were processed for liquid chromatography-mass spectrometry (LCMS) on an Orbitrap Eclipse LCMS system. The resulting mass spectra were input to ProteomeDiscoverer for label-free quantitation (LFQ), with resulting intensity values used for downstream analysis^22,23^. Non-normalized protein intensity data concatenated across samples is listed in Supplementary Table S3.

### Selection of Candidate Surface Markers

Individual sample LFQ intensity values were analyzed within sample groups corresponding to *in vitro* and PDX models, respectively. Within each group, bulk proteins were first checked for the presence of contamination, as indicated by a reference mass spectrometry contaminant database. Additionally, protein intensities were median-normalized to account for differences in intensity distributions between samples. Proteins present in all samples within each sample group – PDX or cell line – were then compared to the Cell Surface Protein Atlas, a mass-spectrometry project to create a compendium of high-confidence cell surface proteins^24^.

### Flow Cytometry Collection and Analysis

Flow cytometry was conducted for a variety of MPNST-derived cell lines using a single-channel approach. Cells were stained with antibodies corresponding to surface targets of interest. Each antibody was conjugated to phycoerythrin (PE) dye and was used at a ratio of 5 µL to 1 million cells. Cell viability was measured using Zombie NIR™ Fixable Viability dye. To control for nonspecific binding, each experiment was compared to an isotype control conjugated to PE. All experiments were performed on a Beckman Coulter CytoFlex 5, and analysis was performed in FlowJo v10.10.

### Quantification of Surface Antigen Density

BD Phycoerythrin QuantiBrite™ beads were used to convert PE fluorescence intensity values to the number of surface antigens on each cell. A single sample of beads was run in parallel to assayed MPNST cell lines, generating 4 distinct peaks of PE-based fluorescence. The median fluorescence intensity of each peak was plotted against the number of PE molecules conjugated to each bead within its tier, both on a log scale. The line of best fit for this relationship was then used to convert median fluorescence intensity for each sample into a median PE molecule number per cell. Each median PE molecule number was then divided by the number of PE molecules per antibody to derive a median antigen density per cell.

### Drug Efficacy Studies and Analysis

Antibody-drug conjugates were assessed for therapeutic efficacy using a medium-throughput dose-response experiment. In brief, cells were plated in 384-well plates on Day 0. On Day 1, drugs were dispensed into wells according to plate design, with each drug-concentration combination being tested in quadruplicate. On Day 5, each plate was assessed for cell viability with Alamar Blue treatment at a 1:10 V/V ratio for 4 hours, then measured for absorbance. Each drug concentration well was shifted by an average intensity of 8 media-only wells per plate, with drug viability calculated as the ratio of the drug-concentration well shifted intensity to the average shifted intensity of 8 wells with zero drug. Data were analyzed and plotted using the R package *drc* with IC_50_ values obtained using a 4-parameter log-logistic function^25^.

### Sample Preparation and RNAseq

Cell pellets of 3-4 million cells were collected. RNA was extracted using the RNeasy Mini Kit (Qiagen) and sent to the University of Minnesota Genomics Center. Sequencing was performed on a NovaSeq 6000 (Illumina), generating an average of 20 million reads/sample with 150 bp paired end reads.

### RNAseq Analysis

RNAseq reads were trimmed and quality filtered using Trimmomatic, then aligned to the human transcriptome by kallisto^26,27^. Before differential expression, transcripts per million (TPM) values were summed over genes and exported to a gene count table using tximport^28^. Differential expression analysis was performed using DESeq2, with a post-analysis filtering step excluding genes with absolute-value log_2_ fold change below 1, and false discovery rate (FDR) above 0.05^29^.

### Statistical Analysis

All statistical analysis was done with the R computing project. Scripts are provided for each analysis for reproducibility. Gene set enrichment analysis (GSEA) and pathway analysis were performed as previously described with the R package *clusterProfiler*^30,31^.

## Results

### Establishing Isogenic MPNST Lines with Controlled PRC2 Activity

To identify molecular targets for MPNST, we isolated proteins present on the cell surface and then identified and quantified these proteins using liquid chromatography with tandem mass spectrometry (LC-MS/MS). The first goal of this approach was to understand the impact of PRC2 activity loss on the cell surface proteome in MPNST. To do so, we generated a model human MPNST cell line with doxycycline-inducible PRC2 activity. S462-TY cells were transduced sequentially with an rtTA cassette followed by a *SUZ12* cassette with a tetracycline regulatory element (TRE)^32,33^. Treatment of these S462-TY “addback” lines with doxycycline for seven days led to complete restoration of H3K27me3 via Western blotting (Supplemental Figure 2). Additionally, RNAseq performed on addback lines with doxycycline and with a DMSO control showed significant degrees of differential expression between S462-TY with PRC2 activity restored and S462-TY lacking PRC2 activity. To understand changes in the surface proteome following PRC2 inactivation, a complementary model to this addback was generated by treating MPNST007, an MPNST-derived cell line with PRC2 activity intact, with an EZH2 inhibitor^34,35^.

### The PRC2 Controls the Abundance of a Distinct Set of Cell Surface Proteins in MPNST Cells

Following the generation of the PRC2-controlled samples of S462TY and MPNST007, each was assayed for its cell surface proteome. Samples were generated for each line with and without PRC2 activity. Comparing the surface protein intensity profiles of samples with PRC2 activity to those without it yields a range of proteins that are suppressed by PRC2 activity and thus are more highly abundant in samples with inactive PRC2 (Figure 1a,b). Many more proteins were preferentially abundant in the PRC2 inactive state than in the active state. This result is consistent with the observed evidence at the mRNA transcript level; that is, the lack of PRC2 activity leads to increased transcription of a wide array of genes, much more so than concomitant mRNA expression decreases^11,36,37^. Comparing the differentially abundant proteins that arose in S462TY and MPNST007 yielded a set of 46 proteins that were consistently more abundant when PRC2 is inactive. This includes receptor tyrosine kinases implicated in cancer, such as ERBB3 and ROR2, as well as proteins involved in cell migration and invasion, such as ITGA4, PLXNA2, and PCDH1. S462TY and MPNST007 exhibited similar effects following PRC2 activity modulation in that PRC2 loss caused surface protein changes. But, there was only a small overlap between the two models in surface proteins whose abundance was significantly higher in the PRC2-inactive set. Pathway analysis was run on both S462TY and MPNST007 independently, assaying the proteins with higher abundance in the PRC2-inactive state. When looking at Hallmark pathways enriched from these protein sets, there are several pathways with overlap^38^. Specifically, of the top ten enriched pathways identified, the pathways of epithelial-mesenchymal transition, interferon alpha response, and fatty acid metabolism are heavily enriched. This result suggests a common role in PRC2 controlling the abundance of surface proteins involved in cell identity and differentiation, immune signaling, and tumor metabolism.

**Figure 1:**
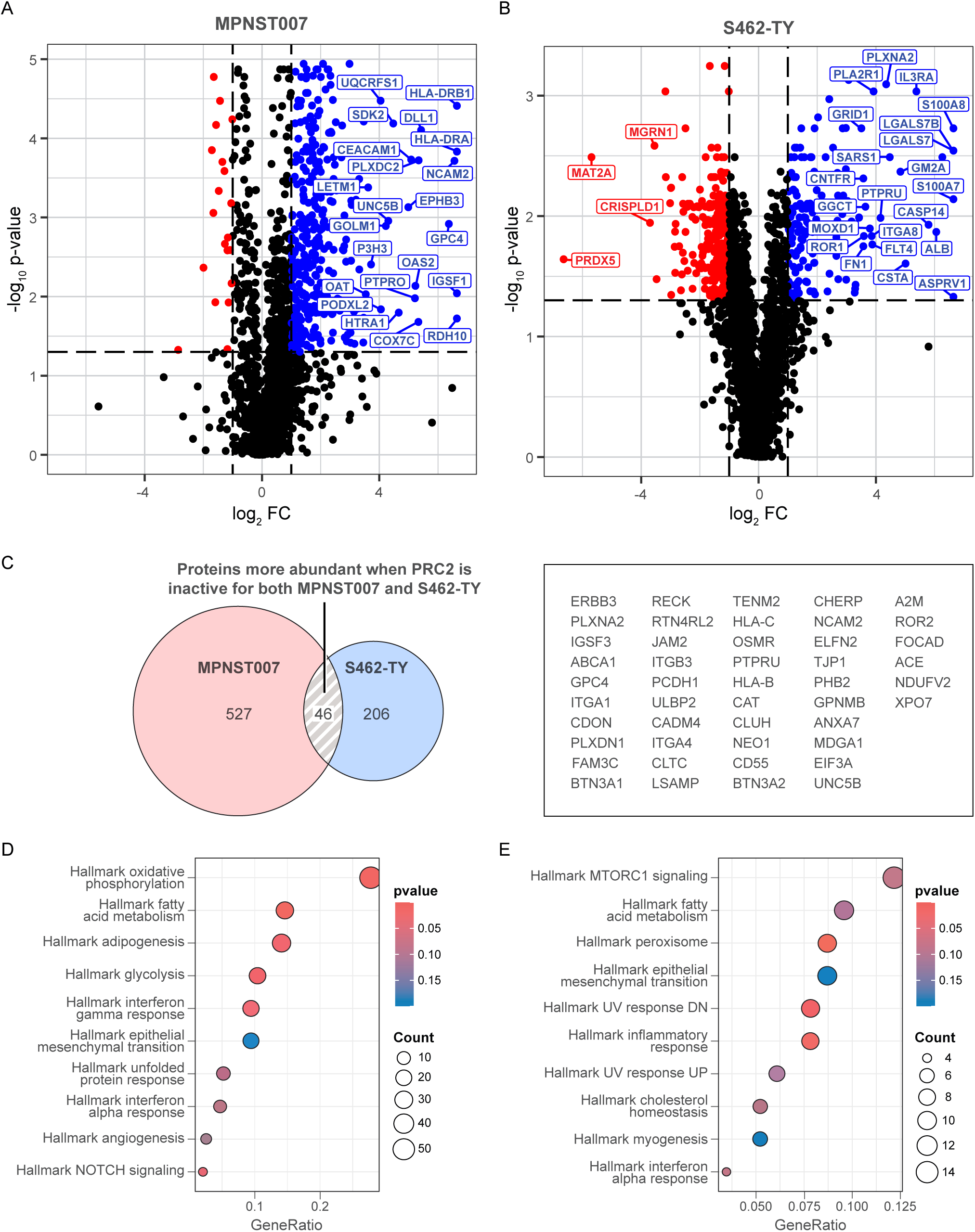
The PRC2-dependent surface proteome. Volcano plots show the most differentially abundant surface proteins for MPNST007 (A) and S462TY (B), both comparing proteins more present when PRC2 is active (blue) against proteins more present when PRC2 is inactive (red). A Venn diagram shows the intersection of lists of proteins more abundant when PRC2 is inactive for both samples, along with a list of those common proteins (C). Pathway dotplots show the enriched pathways from MPNST007 (D) and S462TY (E) PRC2-inactive-up proteins. Despite a relatively shallow intersection of individual proteins, both samples engage similar pathways on the cell surface when PRC2 is inactive.

### mRNA Abundance Poorly Predicts Surface Protein Abundance and the PRC2 Modulates a Subset of Cell Surface Proteins Independently of Changes in mRNA Abundance

Cell surface capture studies were expanded to a wider variety of available tumor-derived cell lines (n = 5) and PDX (n = 7). To evaluate the efficacy and reproducibility of the surface capture approach, a comparison of the resulting mass spectra data to RNAseq run on matched samples was done to understand how well the abundance of RNA transcripts correlated to the abundance of surface proteins. Generally, across several MPNST-derived cell models, RNA abundance (transcripts per million) did not effectively correlate in any direction with surface protein abundance (Figure 2a). However, if low-abundance RNA values are removed, the correlation does improve slightly, suggesting that high variability in lowly expressed RNAs contributes to this. Finally, when binning surface proteins into deciles and comparing RNA expression between deciles, there is no consistent correlation between surface protein abundance decile and RNA expression, suggesting that RNA transcript abundance lacks predictive power to accurately determine how abundant a protein is on the cell surface, even when controlling for different sample intensities. For the most abundant surface protein decile, Pearson correlation between surface abundance and RNA expression rises to 0.12 (p < 0.001), implying that there is a positive relationship between the two. However, the magnitude of this relationship is too small to utilize RNA expression in place of surface protein measurements.

**Figure 2:**
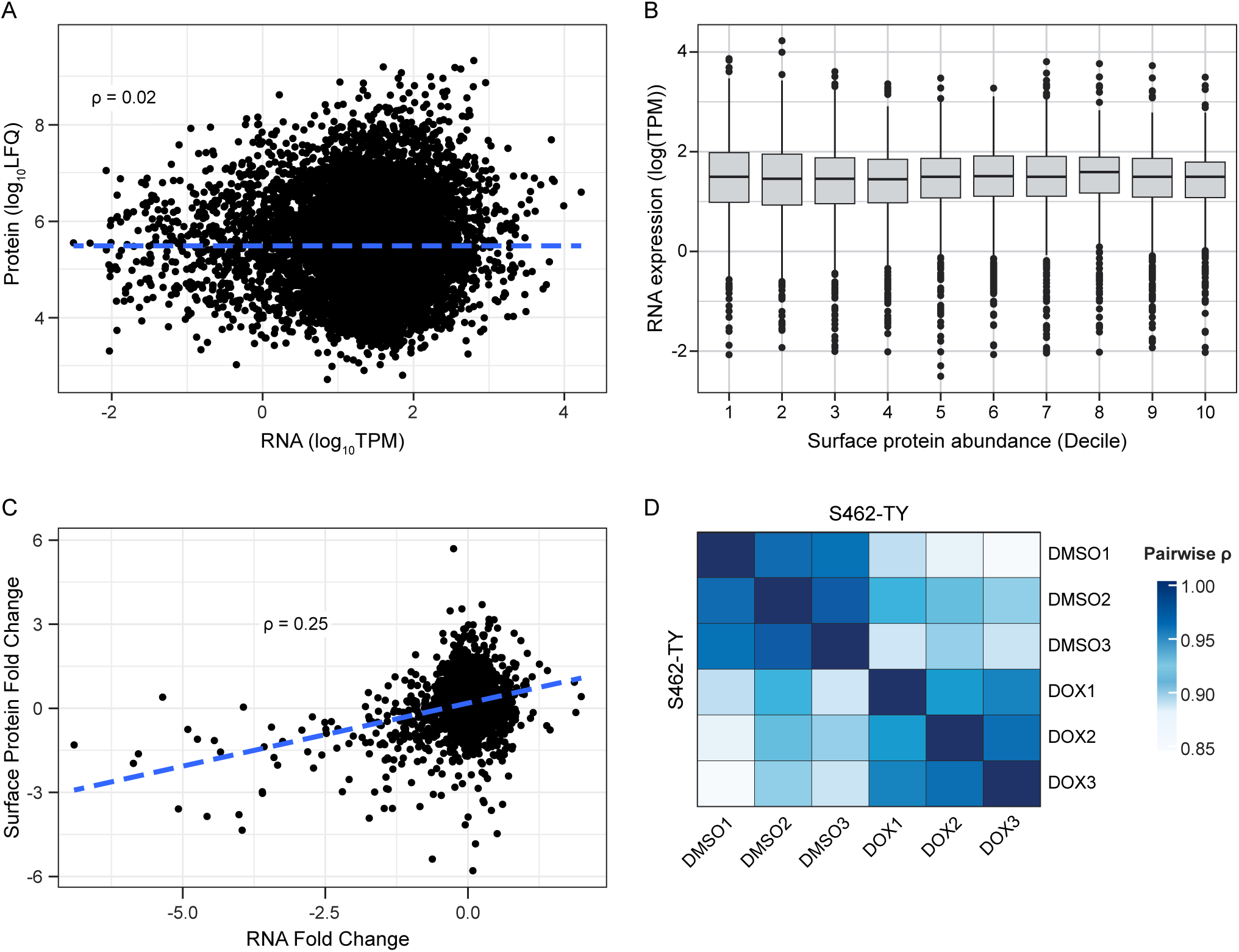
Protein and RNA abundances (A) and experimental fold change values (B) are shown for multiple samples. In general, there is little correlation between a gene’s expression level and its surface abundance. However, changes in abundance at the expression level are positively correlated with changes in surface abundance. Finally, a heatmap (C) shows correlation between technical replicates in their surface proteome abundances. Samples (D) of S462TY with (DOX) and without (DMSO) PRC2 activity are plotted by pairwise sample correlation. Samples within the same group exhibit markedly higher correlation than samples of the opposite group.

Next, changes in transcript level were compared to changes in surface protein in the context of reintroducing PRC2 activity (Figure 2a). When comparing PRC2-inactive samples to PRC2-active samples, fold changes at the RNA and protein level showed a positive correlation (p < 0.05), with changes in RNA abundance tracking with changes in protein abundance across this experiment (Figure 2b). The samples were run as technical replicates, and surface protein abundances were compared within each condition. The mass spectra profiles of these samples correlated highly within their own experimental conditions, with markedly lower correlation to samples of the other experimental condition (Figure 2d). It should be noted that samples across both conditions are isogenic, thus explaining the high correlation regardless of treatment.

Altogether, these results suggest that bulk RNAseq may be able to identify surface proteins with large abundance changes between experimental conditions, but are likely unable to determine the effect size of abundance change. Considering that this is despite controlled experiment conditions using isogenic cell lines, we can infer that real-world RNAseq data is likely to perform worse given the additional biological variation between samples. Additionally, it holds little to no predictive power in determining surface protein abundance from measurements of expression such as TPM, making it a poor proxy measurement in place of surface proteomics.

### MPNST-derived Cell Line and PDX Models Define a Consensus MPNST Surfaceome

Beyond modeling the direct effects of PRC2 activity loss on the cell surface proteome, we also sought to identify a host of proteins consistently expressed on the MPNST cell surface. To do this, data from two sample types were used: PDX and tumor-derived cell lines. For the PDX samples, tumors were obtained from mice and then processed to first homogenize the tumor into a slurry of cells, then separate human tumor cells with magnetic bead separation, depleting both dead and non-human cells for the final PDX sample. Each group was mined for proteins that were detected across each sample in the respective group. Additionally, within each group, proteins that were expressed at less than the group-wise median intensity were discarded. Finally, proteins were compared to a database of proteins that have shown considerable evidence of surface localization^24^.

This workflow yielded 99 proteins that are within the top half of expression in both PDX and cell line models of MPNST (Figure 3). Many of these proteins have been identified across various cancer types as oncogenic drivers, such as PTK7, EGFR, and HER2^39^. An evaluation of mRNA expression for these protein products in human tibial nerve, MPNST, and other precursor neurofibromatosis lesions showed two primary clusters of proteins whose mRNA expression was notably elevated in MPNST and precursor lesions compared to tibial nerve (Supplemental Figure 3). Curiously, before the final narrowing of protein selection to those with abundant cell surface evidence, there was a large amount of divergence between PDX and cell line models in the most highly abundant proteins. Many of these divergent proteins were found exclusively within PDX samples and have little evidence of surface localization, or they were known mass spectrometry contaminants, such as HSPA5, HSPA9, and KRT1^40^. Therefore, it was determined that the most likely cause of these proteins’ appearance in this data is from stromal cells within PDX samples that were not completely removed during tumor dissociation and negative-selection-based removal of mouse cells and dead cells. However, research has implicated that each of the example proteins named above – HSPA5, HSPA9, and KRT1 – potentially localizes to the cell surface, especially upon dysregulation in multiple malignancies. Thus, it is possible these divergent proteins represent biological distinctions between the PDX and *in vitro* cell models^41–43^. Regardless, the final list of proteins provided a robust sample of potential protein targets, a subset of which were chosen to investigate further.

**Figure 3:**
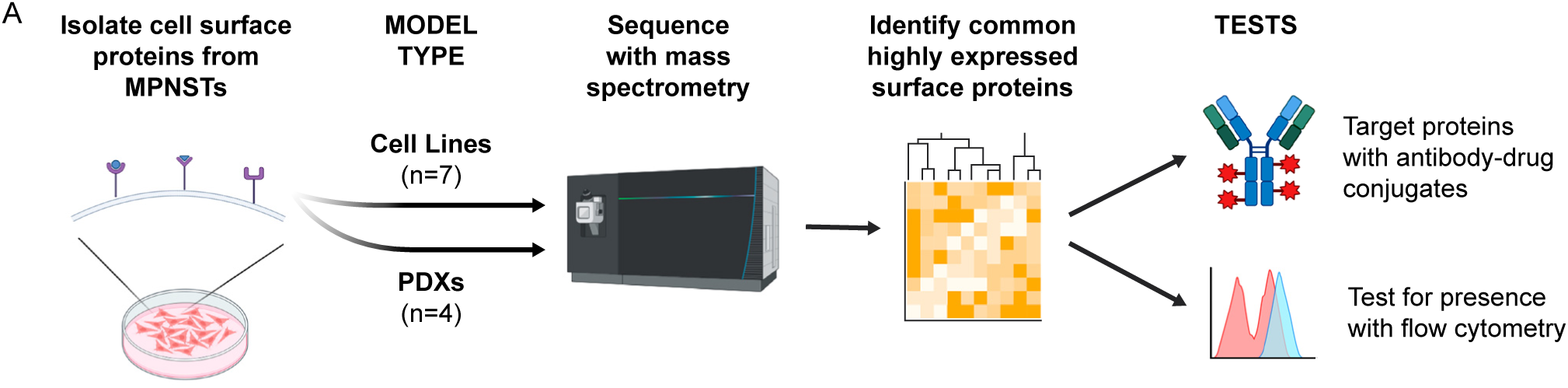
A workflow schematic shows the overview of the cell surface proteome identification procedure. Two distinct model types were used in the form of patient-derived xenografts (PDXs) and cell lines derived from MPNSTs. Upon generating protein spectra, commonalities in protein expression were used to find highly abundant surface proteins, which were then tested by targeting with antibody-drug conjugates and using flow cytometry to confirm their presence on the cell surface.

**Figure 4:**
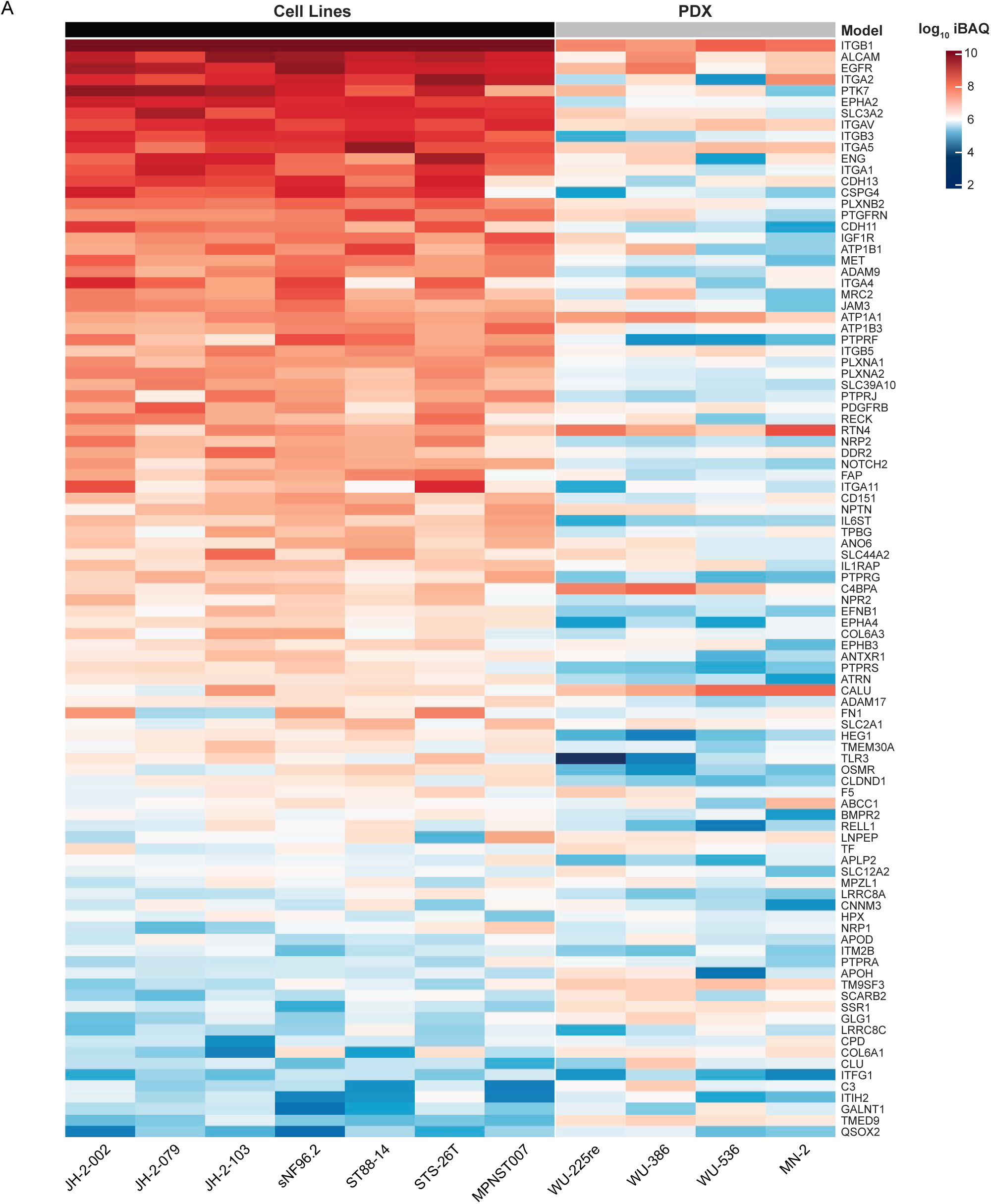
Abundances of surface proteins uniformly expressed across various MPNST models are shown in a heatmap. Each of these 101 proteins is expressed in each of the 11 samples shown. Individual cells shown log10 label-free quantification (LFQ) intensity values. Rows correspond to proteins and columns correspond to the associated sample. Sample types are denoted along the top where gray represents PDX samples and black represents tumor-derived cell lines.

### PTK7 is a Viable MPNST Target

Based on their consistently high abundance across all samples in the final surface proteome dataset, a subset of proteins was selected for further investigation. Specifically, EGFR, HER2, MET, and PTK7 were chosen as targets for validation using flow cytometry and pharmaceutical attack. Surface cell staining detected each of these proteins at levels substantially above isotype controls within each MPNST cell line tested (Figure 5). Notably, the fluorescence for each protein was not directly correlated to the level of intensity in the mass spectrometry data. Of the proteins assayed, PTK7 was expressed consistently at much higher apparent levels than EGFR, MET, and HER2. In each MPNST cell line used, the median fluorescence of PTK7 was roughly an order of magnitude higher than that of the other assayed proteins and another order of magnitude higher than that of the isotype control. Interestingly, two distinct populations were observed when assaying PTK7 in STS-26T, with one group of cells exhibiting fluorescence much closer to the isotype control and another group exhibiting much higher fluorescence than any other protein. This unexpected phenomenon is only observed in PTK7 for this cell line^40–42^. Previous work has shown that the genomic SNP profile of STS-26T is similar to that of cutaneous melanoma; it is also the only cell line in this group derived from a sporadic MPNST^43,44^. These differences could potentially explain the distinct data observed from STS-26T. Despite this, PTK7 is still present at a high level across all assayed lines.

**Figure 5:**
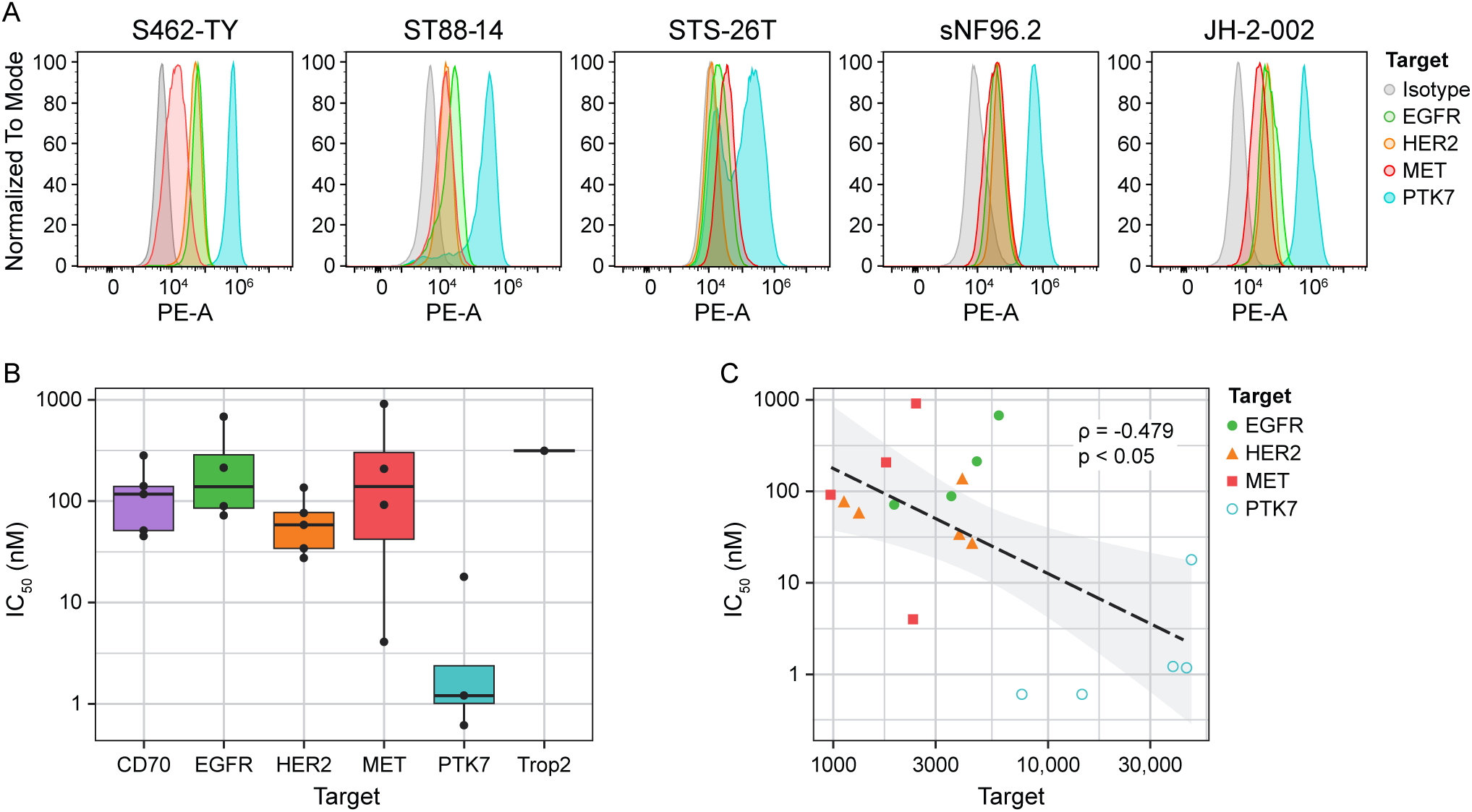
Flow-cytometry and dose-response experiments show both presence of and efficacy targeting surface proteins (A). Antibody-drug conjugates using the same antibody as the corresponding flow cytometry experiments were used to target each cell population for death, with the derived IC50 values for each antigen target plotted in boxplots (B). The correlation between the IC50 values and the surface antigen density – derived from the flow cytometry experiments – is plotted along with a line of best fit (C). A statistically significant negative Spearman’s correlation is observed for this relationship, indicating that the amount of ADC necessary to induce cell death is lower for cells with highly dense antigens on the cell surface.

To understand the potential of targeting the identified surface proteins pharmaceutically, each MPNST cell line was treated with antibody-drug conjugates (ADC) targeted to either EGFR, MET, HER2, CD70, PTK7, or Trop2. Trop2 served as a negative control, given that it was not observed in the mass spectrometry data from either *in vitro* cell lines or PDX models. Identifying IC_50_ values for each cell line-protein target pair shows that each ADC required less than 1 uM to induce a 50% decrease in cell viability (Figure 5). Note that the ADC targeting Trop2 only yielded an IC_50_ value for one cell line, STS-26T, as it did not induce significant cell death in any of the other lines. Of the ADCs used, the one targeting PTK7 consistently exhibited the lowest IC_50_, with one value (assayed in STS-26T) being less than 1 nM. This suggests that, while each surface protein identified in the mass spectrometry data is a useful target for cell death, PTK7 is likely highly abundant on the cell surface of individual MPNST cells and, as such, carries high pharmaceutical relevance as a potential therapeutic target. To further exemplify this potential, we compared mRNA expression of *PTK7* across the above-mentioned tibial nerve, MPNST, and precursor lesion samples and observed that *PTK7* was highly expressed uniquely within the MPNST samples, with gradual increases in expression in the order of tibial nerve, plexiform neurofibromas, ANNUBP, and MPNST (Supplemental Figure 4). This points to PTK7 as both a highly abundant and tumor-specific surface protein, making it an excellent target for therapeutic intervention.

We wished to ascertain whether the physical density of antigens on the cell surface identified from this study could predict the efficacy of antibody-drug conjugates that target these antigens. To do this, the flow cytometry studies were conducted with an additional sample of phycoerythrin (PE) quantification beads, in which known levels of PE conjugated to beads could be used to correlate PE fluorescence measured by the flow cytometer to the number of antigens present on a given cell (Supplemental Figure 5). Calculating the number of antigens on the MPNST cells assayed, we were able to plot the observed antigen density per cell against the IC_50_ values obtained from the ADC studies (Figure 5c). The observed relationship between these values suggests an obvious conclusion in that higher antigen density is predictive of greater ADC efficacy. In this study, specifically, the efficacy of targeting PTK7 is explained by the relatively much higher density of PTK7 on the surface of each MPNST cell.

## Discussion

The results presented here provide a detailed view of the MPNST cell surface proteome. Profiling PRC2-dependent changes to the cell surface proteome offers insight into malignant transformation. Loss of PRC2 likely derepresses a set of cell surface molecules that promote the malignant phenotype of MPNST. Our data indicate that PRC2 loss increases expression of transmembrane receptor tyrosine kinases such as PDGFRA and MET, which are both well-characterized oncogenic drivers in solid tumors^45,46^. Their derepression may enhance proliferative and survival signaling in PRC2-deficient MPNST cells. We also observe elevated expression of integrins such as ITGA1, which mediate extracellular matrix ligands and are known to facilitate tumor invasion and metastasis^47^. This observation is consistent with recent work from Dr. Rebecca Dodd’s laboratory, which demonstrated that PRC2 loss in MPNST cells promotes metastatic behavior *in vivo* via changes in expression for matrix metalloproteases and lysyl oxidases, resulting in tumor extracellular matrix remodeling^48^. We also found derepression of plexins, a family of receptors implicated in cell migration, invasion, and angiogenesis, suggesting that the loss of PRC2 leads to a broader reconfiguration of surface signaling networks that facilitate tumor progression^49^. Together, these findings suggest that PRC2 loss drives MPNST transformation, not only through transcriptional dysregulation but also by altering surface interactions with its microenvironment, enabling malignant behaviors such as growth factor hypersensitivity, invasion, and metastasis. Beyond this, defining the consensus surface proteome of MPNST provides insight into the signaling dynamics driving this disease as well as avenues to disrupt them.

In addition to identifying protein targets for MPNST, this work demonstrates that existing pharmaceutical products can effectively target them. The use of antibody-drug conjugates proves to be remarkably efficient in killing MPNST cells *in vitro* with IC_50_ values in the low nanomolar range, especially when targeting PTK7. These findings support prioritizing surface targets and warrant *in vivo* evaluation of this approach. PTK7 is an emerging, promising candidate for therapeutic targeting across multiple cancer types. Both ADCs and CAR-T therapies have shown promise in models of other malignancies and in clinical trials^38^. Adding to this promise is the fact that all therapeutics used in this study are publicly available and have been used in clinical trials, increasing their overall viability. Finally, *PTK7* mRNA levels are elevated in recurrent MPNST compared to non-recurrent tumors, suggesting additional clinical relevance ^50^.

In addition to PTK7, several other surface proteins identified here have been recognized elsewhere for therapeutic potential in sarcomas. A HER2 antibody-drug conjugate has shown activity in models of desmoplastic small round cell tumors (DSRCT), a soft tissue sarcoma similar to MPNSTs^51^. In MPNST specifically, evaluation of transcriptomic subtypes of MPNST has identified an “immune-deficient” subtype of MPNST characterized by elevated *HER2* expression relative to other MPNST subtypes within the work^50^. Finally, a multi-center study of *HER2* mutation frequency found that nearly 40% of MPNST within the study harbored *HER2* amplification^52^. Each of these data suggests targeting of HER2 could be an effective clinical strategy. Evidence exists for potential MET targeting as well. A study examining responses to a common treatment modality for MPNST, MEK inhibition, noted that MET is commonly upregulated following treatment^53^. In addition, work combining mass spectrometry analysis of MPNST tumors with drug screening identified MET as significantly enriched in local recurrence samples of MPNST, while two MET inhibitors, crizotinib and foretinib, showed standout antitumor activity and were proposed as novel drugs for MPNST treatment^18^.

Specific questions do remain after evaluation of the results shown here. For instance, validation of these surface markers was done exclusively in tumor-derived cell lines due to their ease of handling and quick turnaround time. Further work in *in vivo* models would elucidate the effectiveness of these approaches in more clinically relevant contexts. The relative homogeneity of cell line models is a limitation, as patient lesions exhibit a fair degree of intratumor heterogeneity, tumor microenvironment and surrounding stroma notwithstanding^54,55^. However, evidence from studies of MEK inhibitors in MPNST patients does show that the expression of markers such as EGFR rises after treatment with MEKi, suggesting that combining antibody-drug conjugates with molecular inhibitors might be a worthwhile approach^53^. Additionally, the extent of co-expression among these targets remains unclear. In a scenario in which a subset of tumor cells is highly abundant in PTK7 and a distinct subset are abundant in HER2, it is feasible to imagine a combination of PTK7 and HER2 ADCs producing a highly effective clinical impact and more complete tumor eradication.

Although the work shown here evaluates the abundance of surface proteins in cell lines derived from MPNST and from PDXs, it does not contain proteomic data taken directly from clinical MPNST samples. The samples used here are likely more homogeneous than primary tumors, limiting our understanding of how widespread markers such as PTK7 are expressed within the tumor and surrounding microenvironment and how the abundance of these markers corresponds to patient outcomes. Some work has been done in this respect, namely in defining transcriptomic subtypes of primary MPNST and their associated outcomes and molecular features^50,55^.

Ultimately, these studies comprise a compendium of proteomic and transcriptomic data with associated validation through drug viability screening and flow cytometry. In turn, we hope this data is mined by other groups studying MPNST. Given the clear lack of predictive power in RNAseq in estimating cell surface abundance, we hope that this resource serves as a valuable supplement to studies seeking to identify new molecular targets in MPNST. Additionally, we wish to expand this resource by incorporating additional model systems to increase its comprehensiveness. With this stated, it is important to consider the potential for false positives. The discrepancies observed between cell-line- and PDX-based models illustrate the lack of specificity of this experimental approach. While correlation between technical replicates is quite high, the high sensitivity of mass spectrometry lends itself to the identification of spurious targets. Caution should be taken in assessing the results, specifically in cases where there is a lack of evidence. Use of multiple sample types allows for a more robust comparison, but there remains potential for false positives nonetheless. However, the validation efforts undertaken here show that this method does succeed in its most fundamental aim in identifying consensus surface proteins.

## Authorship statement

Designed the study: C.M.S., L.W., D.A.L. Advised the study: C.M.S., D.A.L. Acquired funding: D.A.L. Acquired and generated cell lines and PDX: C.M.S., Z.S. Performed experiments and analyzed data: L.W., C.M.S. Performed bioinformatic and statistical analysis: C.M.S. Wrote the manuscript: C.M.S., D.A.L.

## Ethics Statement

Protocols used during the course of this study were approved by the Ethics Committee of the University of Minnesota.

## Acknowledgments

The authors would like to thank Dr Nancy Ratner for providing MPNST cell lines, Dr Christine Pratilas for providing MPNST cell lines and cell lines for PDX engraftment, and Dr Angela Hirbe for providing cell lines for PDX engraftment. We acknowledge Dr Nancy Ratner and Dr Christine Pratilas for import advice and discussion regarding the planning of this work. We acknowledge Alex Larsson for work to main PDX samples and for managerial support for the lab. We thank Eva Mae Baucom for assistance with graphic design. We thank WordMedic for assistance with editing and revision. The use of generative AI and AI-assisted technologies was not used in the preparation or editing of this manuscript and the authors accept full responsibility for the content of this article, including analysis.

## Data Availability

The raw proteomic data is available upon request. Please for inquiry. Transcriptomic data used in this study is available at https://www.ncbi.nlm.nih.gov/geo/query/acc.cgi?acc=GSE263107.

---

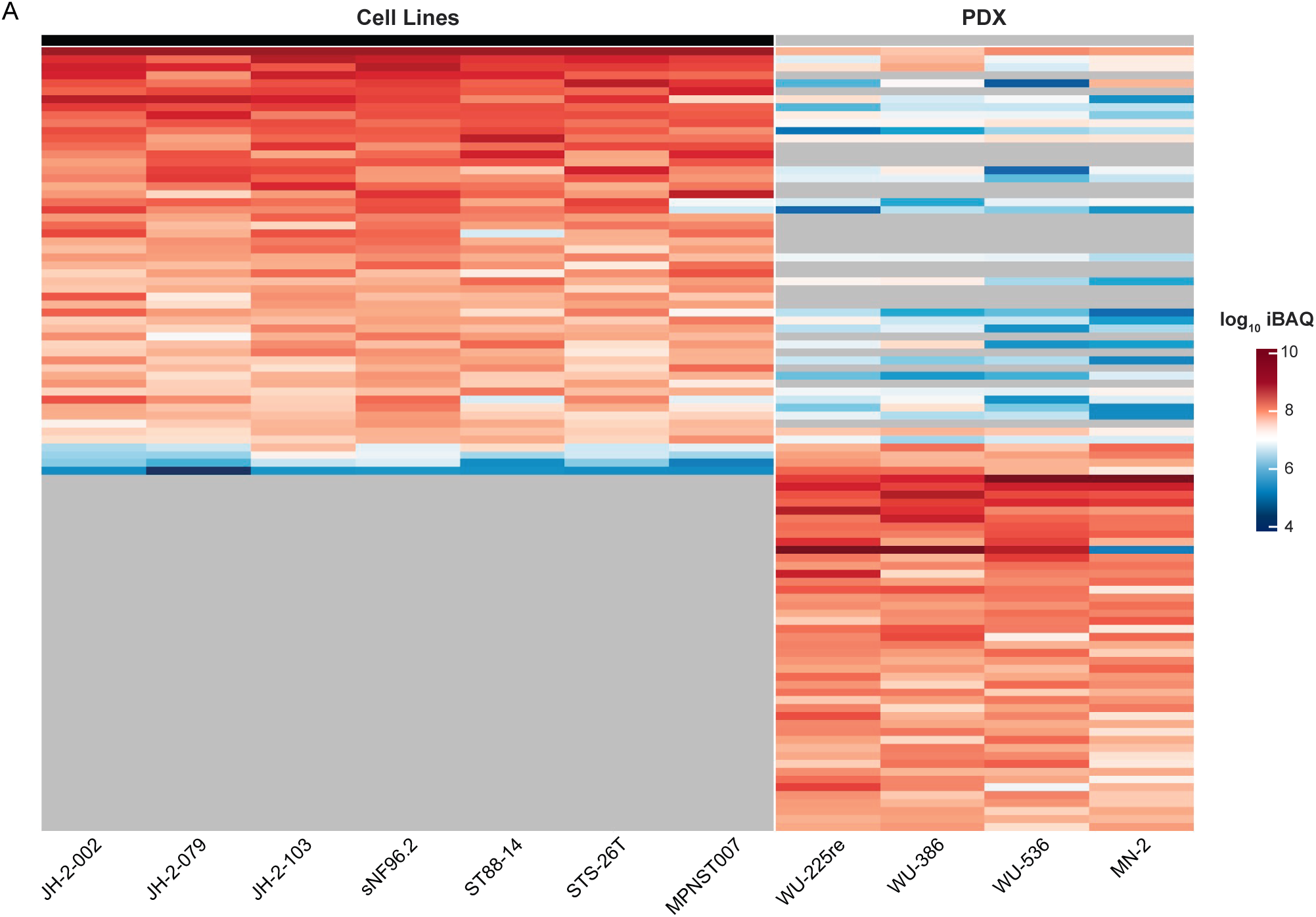

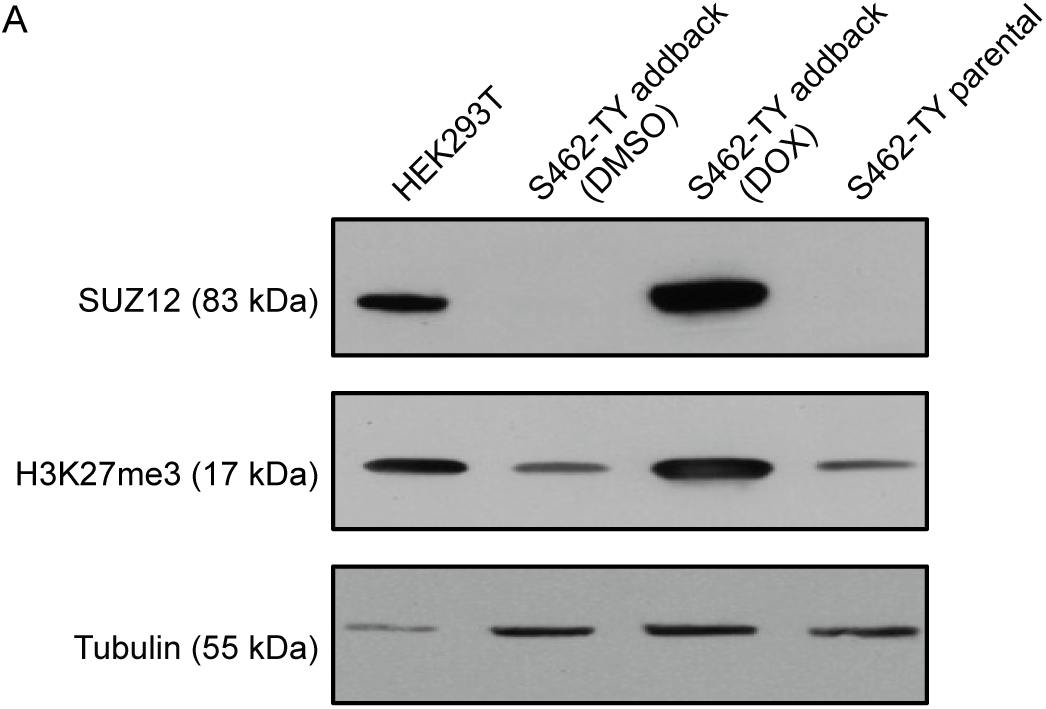

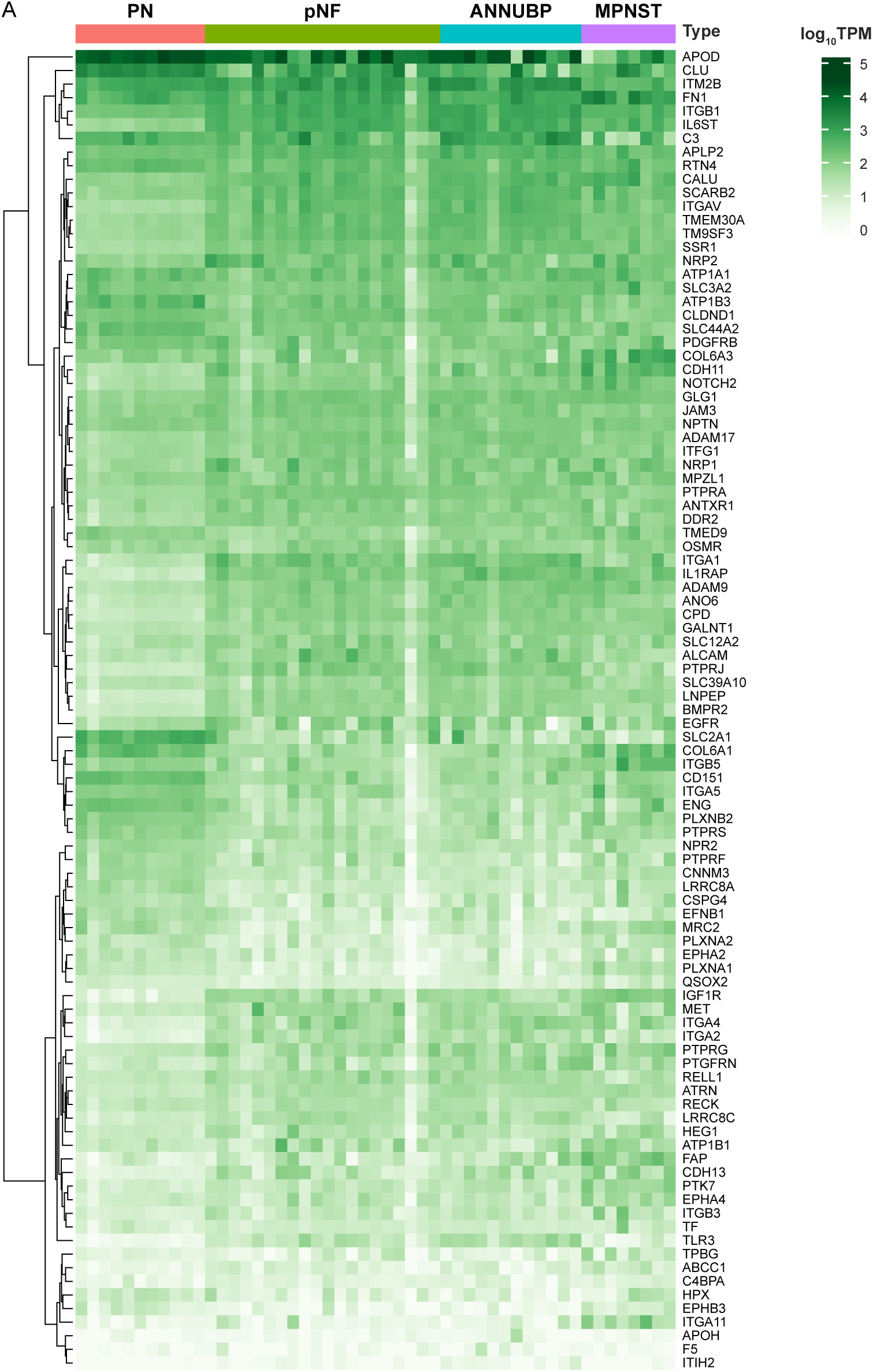

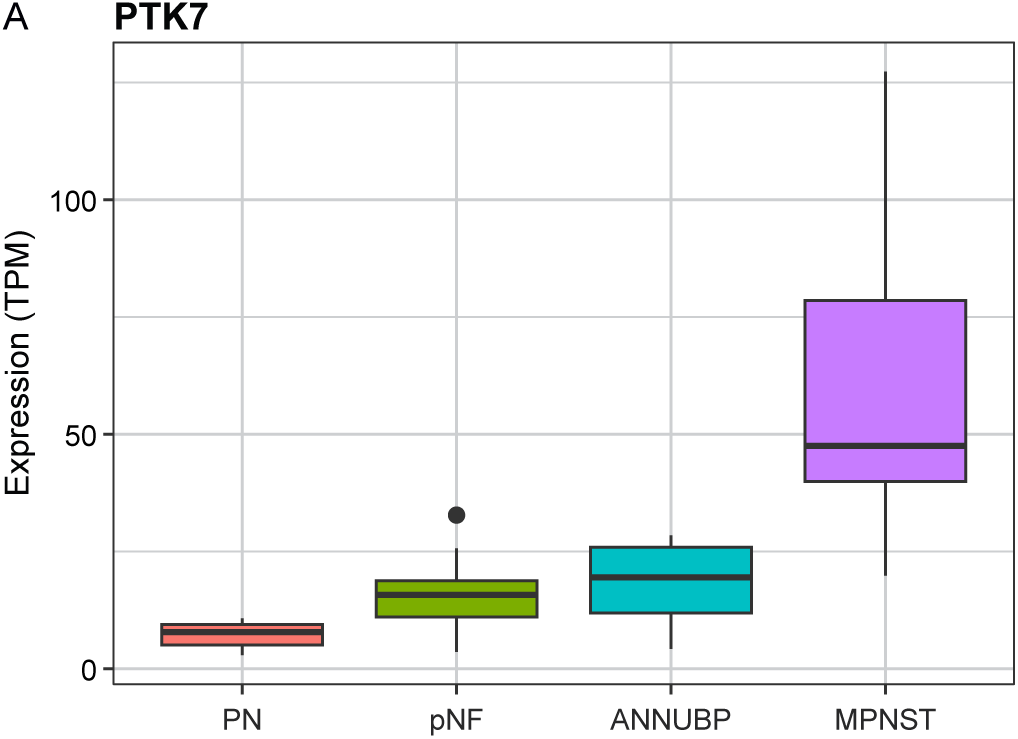

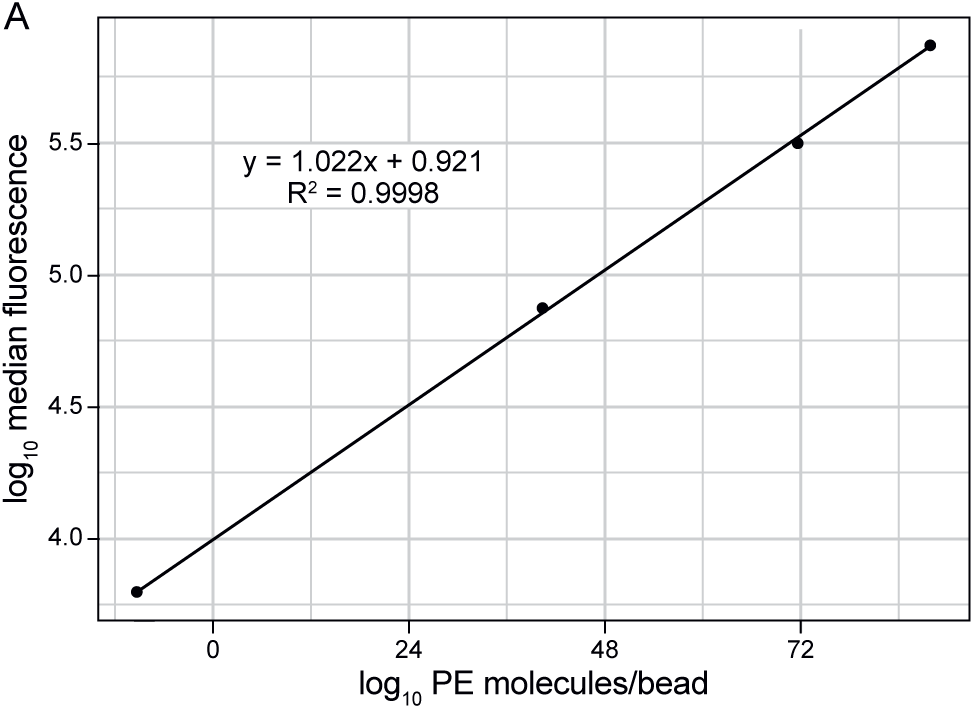

